# Exploring the mechanism of *kaji-ichigoside F1* on bacterial pneumonia based on the network pharmacology and transcriptome

**DOI:** 10.1101/2025.04.17.649383

**Authors:** Junlin Yang, Xi Yang, Liping Wu, Cheng Min

## Abstract

**Objective:** To explore the anti-inflammatory mechanism of *kaji-ichigoside F1* against bacterial pneumonia based on transcriptomics and network pharmacology.

**Method:** Network pharmacological was used to analyse the potential target genes of *kaji-ichigoside F1* action on bacterial pneumonia; molecular docking was used to analyse the docking binding energy of *kaji-ichigoside F1* with key target genes; RAW264.7 macrophages were treated with *klebsiella pneumoniae* fluid and given *kaji-ichigoside F1* intervention, the expression of inflammatory factors IL-1β, IL-6, TNF-α and IL-10 were dectected by qRT-PCR; transcriptomic analysis was performed to obtain differentially expressed genes, and relevant signaling pathways.

**Result:** Network pharmacological analysis showed that the five the key target genes for *kaji-ichigoside F1*-bacterial pneumonia interaction were were TLR4, NFKB1, STAT3, IL1B, and JUN; molecular docking results of *kaji-ichigoside F1* with key target genes showed the docking binding energy ranging from -5.9 to -8.4 kcal/mol; *kaji-ichigoside F1* can reduce the *klebsiella pneumoniae* induced inflammatory response of macrophages, manifested with reducing the mRNA expression of pro-inflammatory factors IL-1β, IL-6 and TNF-α, and increase the mRNA expression of anti-inflammatory factor IL-10; transcriptomics analysis showed that the signaling pathways involved were mainly the TLR signaling pathway and NFKB signaling pathway.

**Conclusion:** This study showed that *kaji-ichigoside F1* alleviates bacterial pneumonia by targeting TLR and NFKB signaling pathways, rebalancing macrophage polarization with suppressing pro-inflammatory cytokines and increasing anti-inflammatory cytokines, highlighting *kaji-ichigoside F1* as a novel agent for combating bacterial infections.

**Graphic abstract:** 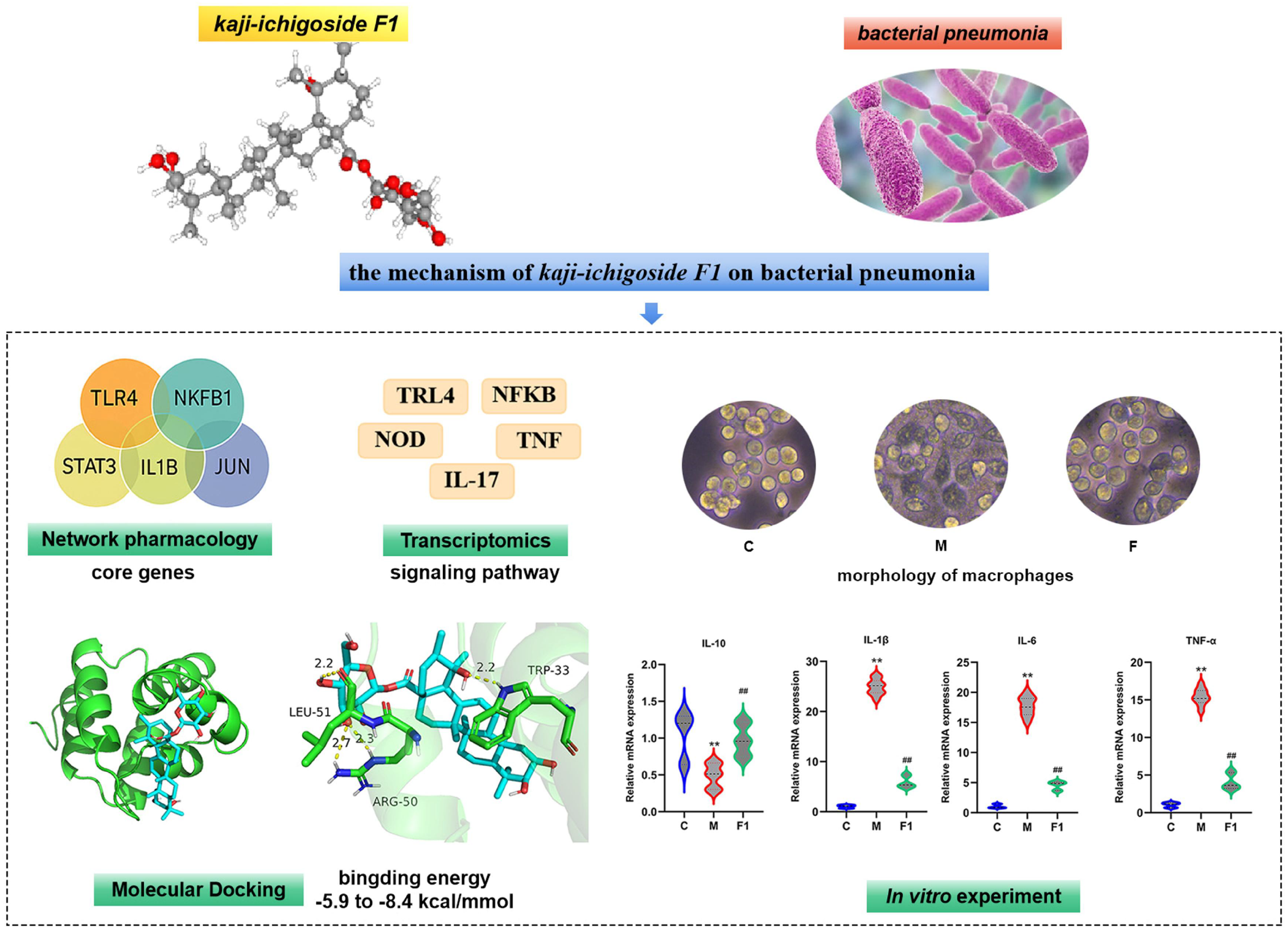

## 1. Introduction

Bacterial pneumonia is one of the most common infectious diseases, the global incidence, morbidity, and mortality of bacterial pneumonia have risen significantly worldwide^[1]^. While antibiotics remain a cornerstone treatment, their overuse has led to an increase in bacterial drug resistance rate year by year, resulting in a significant increase in the difficulty of treatment and posing a serious challenge to global public health^[2]^. In recent years, more and more researchers have started to pay attention to the research in the field of “new antibacterial drugs based on natural compounds” under the concept of green pharmaceuticals and safe use of medicines, botanical compounds have emerged as a significant research focus^[3]^.

*Rosa roxburghii* is the fruit of reeling flower of rosaceae family, which is mainly distributed in the southwest of China, among which is the most abundant resource in Guizhou province^[4]^. Phytochemical profiling reveals its bioactive constituents comprise polysaccharide complexes, flavonoid derivatives, superoxide dismutase enzymes, ascorbic acid, and triterpenoid metabolites. Triterpenoids, as ubiquitous phytoconstituents in higher plants, have undergone rigorous scientific scrutiny for their pleiotropic pharmacological actions, confirming their inflammatory pathway modulation and anti-oxidative properties^[5-8]^. Notably, *kaji-ichigoside F1* emerges as the principal constituent among triterpenoid isolates from *rosa roxburghii*, constituting the predominant pharmacologically active fraction.

Macrophages, central orchestrators of pulmonary immunity, dynamically shift between pro-inflammatory M1 and anti-inflammatory M2 phenotypes during infection^[9]^. Excessive M1 polarization driven by Toll-like receptor (TLR) signaling exacerbates tissue damage through NFKB mediated cytokine storms (IL-6, IL-1β, TNF-α), while impaired M2 resolution perpetuates chronic inflammation^[10]^. Natural compounds capable of rebalancing this polarization, such as ursolic acid in *rosmarinus officinalis* have demonstrated therapeutic efficacy in pneumonia models^[11]^. However, despite its antimicrobial promise, the mechanism by which *kaji-ichigoside F1* alleviates bacterial pneumonia, particularly its immunomodulatory effects on macrophage polarization, remains unexplored.

To further investigate the possible pathways and targets of *kaji-ichigoside F1* for the treatment of bacterial pneumonia, we integrated network pharmacology, transcriptomics, and molecular docking to explore the role and mechanism of *kaji-ichigoside F1* in ameliorating bacterial pneumonia, and to develop novel antimicrobial drugs based on natural compounds.

## 2. Materials and Methods

### 2.1. Network pharmacology analysis

#### 2.1.1. Potential targets of *kaji-ichigoside F1* and bacterial pneumonia

The Pubchem database (https://pubchem.ncbi.nlm.nih.gov/) was searched for *kaji-ichigoside F1*, and the SMILE numbers were entered into the Swiss Target Prediction database, the SEA database, and the Superpred database to summarize and removed duplicates to obtain targets. Using Gene Card human gene database and OMIM database (https://omim.org/), we searched with “bacterial pneumonia” as the search term, set the species as “Homo sapiens”, summarized the disease targets genes and removed the duplicates.

#### 2.1.2. Protein-Protein Interaction (PPI) Network Construction

The targets shared by *kaji-ichigoside F1* and bacterial pneumonia were entered into the STRING database (https://string-db.org/cgi/input.pl), and the PPI network was constructed by setting the species as “Homo sapiens”. The tsv. files obtained from the STRING database were imported into cytoscape 3.7.2 software to analyze the topology of the PPI network, and the key target genes for *kaji-ichigoside F1*-bacterial pneumonia interaction were screened out according to the degree value ranking. In order to better understand the complex interactions between *kaji-ichigoside F1* and their corresponding targets, *kaji-ichigoside F1*-target network diagrams were constructed based on the incorporated compounds and targets, and imported into cytoscape 3.7.2 software for network mapping.

#### 2.1.3. Enrichment Analysis

Gene Ontology (GO) enrichment and Kyoto Encyclopedia of Genes and Genomes (KEGG) pathway enrichment were conducted using DAVID database (https://david.ncifcrf.gov/). GO Enrichment included biological process (BP), cellular component (CC) and molecular function (MF). *P* < 0.05 and FDR < 0.05 were recognized as significant of GO Enrichment and KEGG Enrichment. Visualization and analysis of the results using microbiometrics (http://www.bioinformatics.com.com.cn/).

### 2.2. Molecular docking analysis

Compound name, molecular weight and 3D structure of *kaji-ichigoside F1* were determined from the PubChem database, and the 3D structure corresponding to the active ingredient was downloaded from the RCSB PDB database (http://www.rcsb.org/). Then, using autodocktools software (AutoDock Vina), the ligands and proteins required for molecular docking were prepared, and for the target proteins, their crystal structures were removed from water molecules, hydrogenated, modified with amino acids, optimized for energy and adjusted for force-field parameters, and after that, low-energy conformations of the ligand structures were satisfied, and a library of small-molecule compounds was constructed at the same time. Finally, these key target structures are molecularly docked with the key active ingredient structures, and their score value represents the score of docking between the two, the smaller the score value, the better the result.

### 2.3. Experimental vertification *in vitro*

#### 2.3.1. Cell culture and treatment

In this study, RAW264.7 macrophage were used to construct an cell model of *klebsiella pneumoniae* infection and a *kaji-ichigoside F1* intervention. The liquid concentration of *klebsiella pneumoniae* was adjusted to 0.5 McLeod concentration with a turbidimeter, which was equivalent to 1.5×10^8^cfu/mL, then, the bacterial solution was diluted three times at the ratio of 1:10 to obtain the bacterial solution with the number of bacteria in (1-2)×10^7^, 10^6^ and 10^5^cfu/mL. RAW264.7 cells were treat with 10^7^, 10^6^ and 10^5^ cfu/mL bacterial solution for 4 h. *Kaji-ichigoside F1* were purchased from KKL Med Inc. RAW264.7 cells were treat with 1, 5 and 25 umol/L *kaji-ichigoside F1* for 4 h. The CCK8 method was used to detected the *klebsiella pneumoniae* and *kaji-ichigoside F1* on the viability of RAW264.7 cells. Based on the cell viability test results, the concentration 10^7^ cfu/mL *klebsiella pneumoniae* bacterial solution for 4 h were used to construct model group (M group) and 107 cfu/mL *klebsiella pneumoniae* bacterial solution and 5 umol/L *kaji-ichigoside F1* acted simultaneously on the cells for 4 h as the *kaji-ichigoside F1* intervention group (F1 group), and a blank control group (C group) was set up.

#### 2.3.2. Transcriptomics sequencing

We extracted RNA from the cells in each group according to the instructions of RNA extraction kit, and three wells of each group were combined into one sample. cDNA was amplified by Ovation RNA-Seq System, cDNA library was constructed by SPRI works Fragment Library System II, and the extracted RNA solution was entrusted to Shanghai Sangong Bioinformatics Technology Co. Ltd. for transcriptomic sequencing. The differentially expressed genes in the control group vs. model group and model group vs. intervention group were screened by |log_2_FC|>1 and *P*<0.05, and the differentially expressed genes were enriched by GO and KEGG pathways.

#### 2.3.3. Quantitative Real-Time PCR

According to the manufacturer’s instructions, total RNA was extracted from different groups of samples separately. Using reverse transcription PCR, RNA was first reverse transcribed to obtain cDNA. cDNA was then used as a template for PCR amplification: pre-denaturation at 95 °C for 10 min, denaturation at 95 °C for 20 s, annealing at 60°C for 30 s, and 40 cycles were performed. The relative expression of IL-10、IL-1β、IL-6 and TNF-α mRNA in each group was calculated using the 2^-ΔΔCt^ relative quantification method with GAPDH as an internal reference. Table 1 shows the primer sequences of all genes which were synthesized by Takara Biomedical Technology Co., Ltd.

**Table 1.**
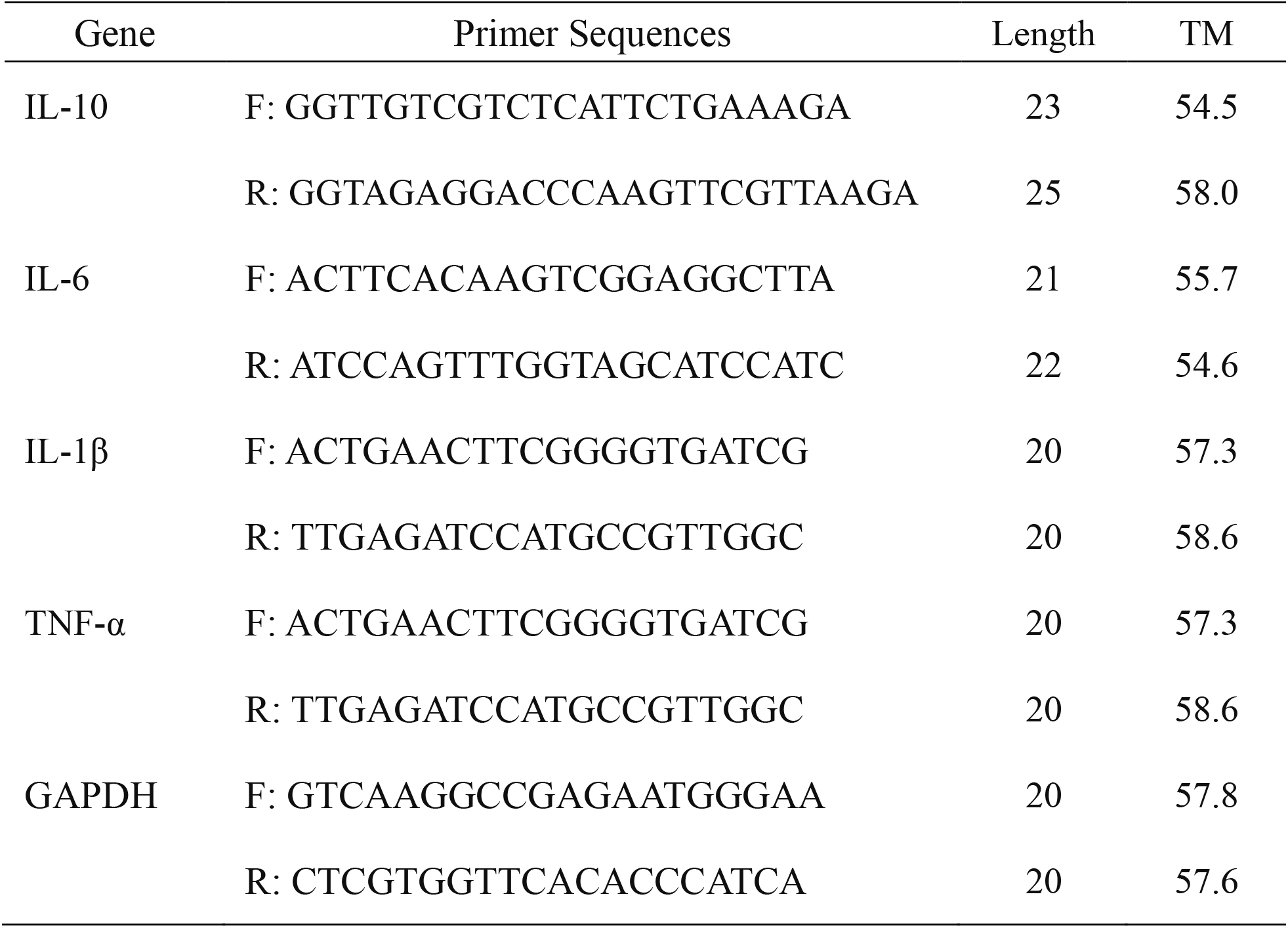
Primer sequence and amplification length of destination fragment.

### 2.4. Statistical analysis

Statistical analyses were carried out with IBM SPSS Statistic23.0 software. Data were presented as the mean ± standard deviation (SD). The T test was used for pairwise comparison, *P* < 0.05 was considered statistically significant.

## 3. Results

### 3.1. Network pharmacological analysis

#### 3.1.1. Venn diagram

A total of 118 *kaji-ichigoside F1* component targets were retrieved using the database, and 6235 bacterial pneumonia disease targets were retrieved, and the screened drug targets and disease targets were entered into the Wayne Diagramming software Venny 2.1, yielding 47 shared targets as predicted targets for *kaji-ichigoside F1* action on bacterial pneumonia for subsequent analysis, as shown in figure 1A.

**Figure 1.**
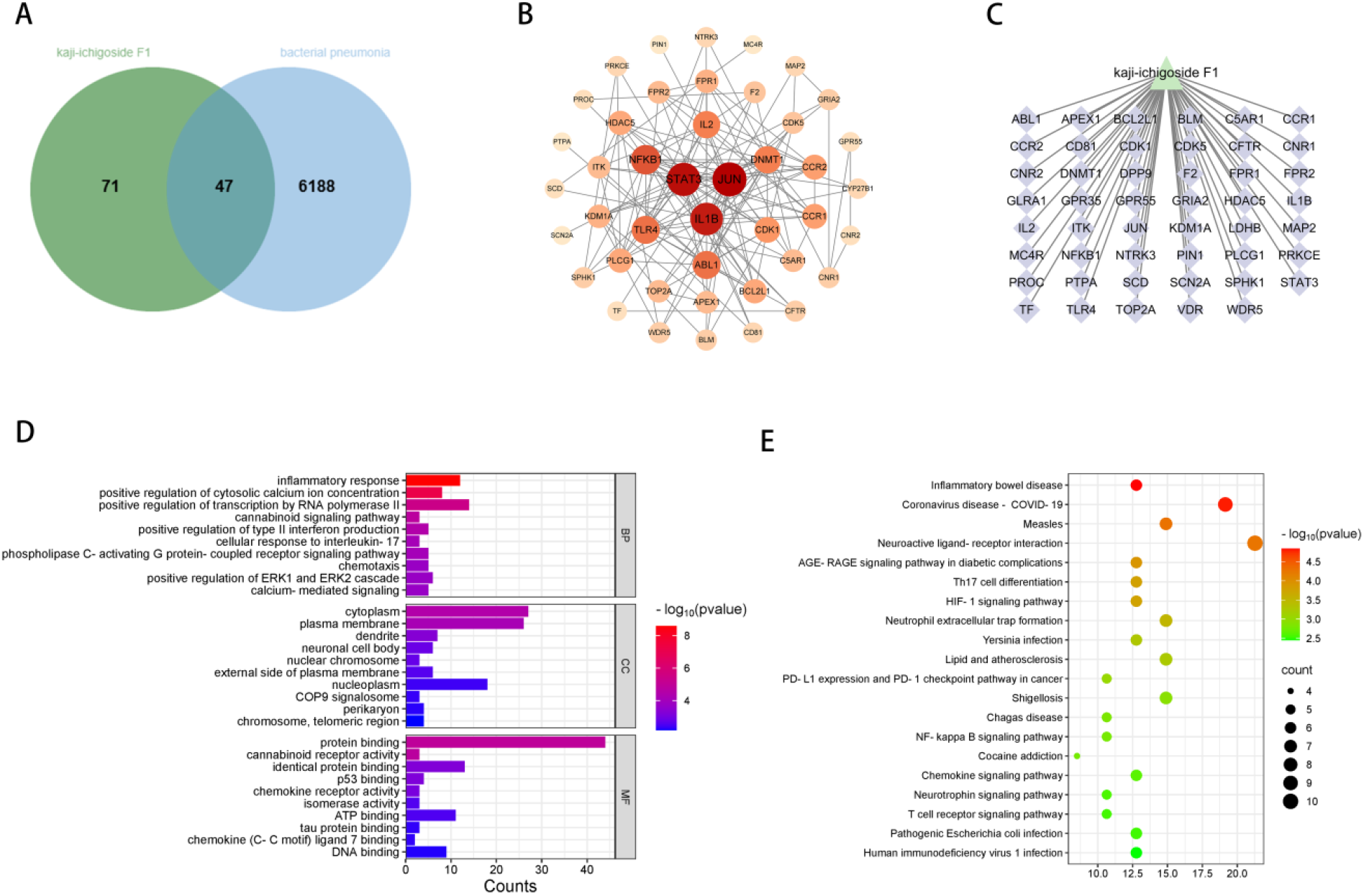
Network pharmacological analysis. A: Venn diagram of *kaji-ichigoside F1* action on bacterial pneumonia; B: PPI network of *kaji-ichigoside F1*-bacterial pneumonia; C: *kaji-ichigoside F1*-target network; D: GO enrichment; E: KEGG enrichment.

#### 3.1.2. PPI network

There were 47 nodes and 142 edges in the PPI network, and the average degree value was 6.17. The PPI network was topologically analyzed, and the key target genes of *kaji-ichigoside F1*-bacterial pneumonia were screened out based on the ranking of the degree value, which were JUN, STAT3, IL1B, NFKB1, and TLR4, with the degree values of 21, 20, 19, 15, and 13, respectively, as shown in figure 1B. The *kaji-ichigoside F1*-target network diagram was constructed based on the included compounds and targets of action, as shown in figure 1C.

#### 3.1.3. GO enrichment analysis

A total of 192 GO entries were enriched. There were 25 CC mainly localized in the cytoplasm, plasma membrane, dendrites, neuronal cell bodies, nuclear chromosomes, outer plasma membrane, nucleoplasm, COP9 signalosome, perinucleosomes, and chromosomal telomeric regions. 129 BP mainly regulating and participating in inflammatory responses, positive regulation of cytoplasmic calcium ion concentration, positive regulation of transcription by RNA polymerase II, cannabinoid signaling pathway, positive regulation of type II interferon production, cellular responses to interleukin-17, phospholipase C-activated G-protein-coupled receptor signaling pathway, chemotaxis, positive regulation of ERK1 and ERK2 cascades and calcium-mediated signaling, among other processes. 38 MF mainly in protein binding, cannabinoid receptor activity, identical protein binding, p53 binding, chemokine receptor, isoenzyme activity, ATP binding, tau protein binding, chemokine (C-C motif) ligand 7 binding, and DNA binding, as shown in figure 1D.

#### 3.1.4. KEGG enrichment analysis

A total of 56 signaling pathways were enriched. The major enrichments were in signaling pathways such as inflammatory bowel disease, coronavirus disease-neococcal pneumonia, measles, neuroactive ligand-receptor interactions, AGE-RAGE signaling pathway in diabetic complications, Th17 cell differentiation, HIF-1 signaling pathway, neutrophil extracellular trap formation and NFKB signaling pathway, as shown in figure 1E.

### 3.2. Molecular docking result

As shown in figure 2, molecular docking results of *kaji-ichigoside F1* with key target genes showed that *kaji-ichigoside F1* formed hydrogen-bonding interactions with VAL-41, MET-20 of the target proteins of IL1B, the docking binding energy was -8.0 kcal/mol. *Kaji-ichigoside F1* formed a hydrogen-bonding interaction with VAL-312, a target protein of JUN, the docking binding energy was -5.9 kcal/mol. *Kaji-ichigoside F1* formed hydrogen-bonding interactions with LEU-51, ARG-50, and TRP-33 of the target proteins of NFKB1, the docking binding energy was -7.4 kcal/mol. *Kaji-ichigoside F1* formed hydrogen-bonding interactions with PRO-333, ALA-250, ARG-350, the target proteins of STAT3, the docking binding energy was -8.4 kcal/mol. *Kaji-ichigoside F1* formed hydrogen-bonding interactions with HIS-426, LYS-402 of the target proteins of TLR4, the docking binding energy was -7.6 kcal/mol.

**Figure 2.**
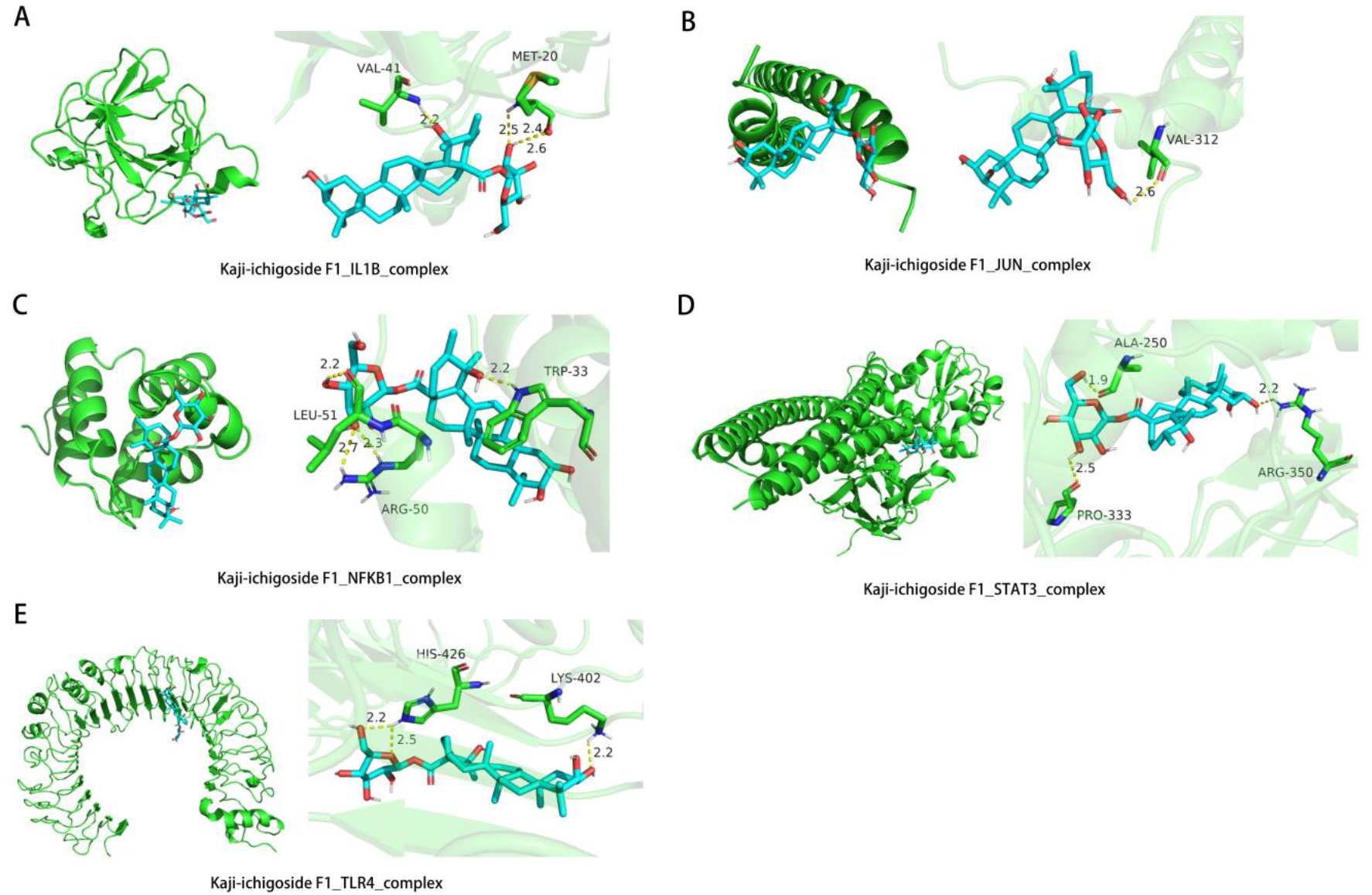
Molecular docking result. A: *kaji-ichigoside F1*-IL1B complex; B: *kaji-ichigoside F1*-JUN complex; C: *kaji-ichigoside F1*-NFKB1 complex; D: *kaji-ichigoside F1*-STAT3 complex; E. *kaji-ichigoside F1*-TLR4 complex.

### 3.3. Results of *in vitro* experiment

According to the result of CCK8, the concentration 10^7^ cfu/mL *klebsiella pneumoniae* bacterial solution for 4h were used to construct model group (M group) and 10^7^ cfu/mL *klebsiella pneumoniae* bacterial solution and 5 umol/L *kaji-ichigoside F1* acted simultaneously on the cells for 4h as the intervention group (F1 group), as shown in figure 3A. In the control group, macrophages have a well-defined circular or oval shape, in the model group, the cells protruded the pseudopodia, became spindle or polygon, the volume increased, the envelope darkened. In the intervention group, the abnormal cell morphology induced by *klebsiella pneumoniae* stimulation was improved to different degrees, the cell morphology was similar to round, as shown in figure 3B. The results of qRT-PCR showed that *kaji-ichigoside F1* as a drug intervention can reduce the mRNA expression of pro-inflammatory factors IL-1β, IL-6 and TNF-α, and increase the mRNA expression of anti-inflammatory factor IL-10 in *klebsiella pneumoniae* induced RAW264.7 cells, suggesting that *kaji-ichigoside F1* can reduce the *klebsiella pneumoniae* induced inflammatory response of macrophages, as shown in figure 3C.

**Figure 3.**
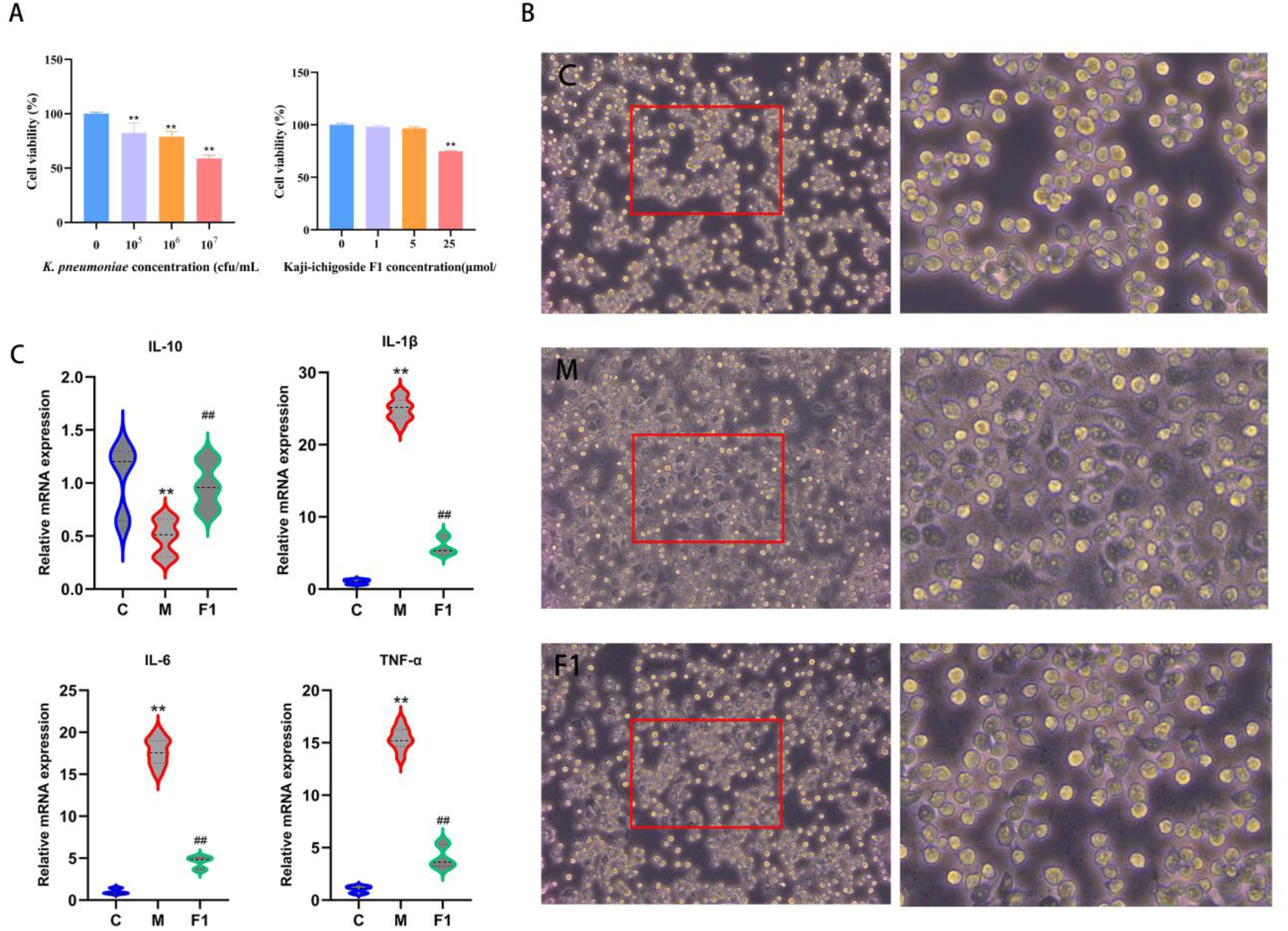
Results of in vitro experiment. A: CCK8 results of *klebsiella pneumoniae* and *kaji-ichigoside F1* on the viability of RAW264.7 cells; B: Macrophage morphological changes in control group, model group, and intervention group (left 10x magnification, right 40x magnification); C: The mRNA expression of inflammatory factors, the data are shown as the mean±SD, compared with the control group, \*\**P*<0.01; compared with the model group, ^##^*P*<0.01.

### 3.4. RNA sequencing results

#### 3.4.1. analysis of differentially expressed genes

In order to further explore the biological characteristics of RAW264.7 macrophages affected by *klebsiella pneumoniae* and *kaji-ichigoside F1* intervention, mRNA transcripome sequencing was performed on macrophages in model group, intervention group and control group. As shown in figure 4A, 850 up-regulated genes and 2112 down-regulated genes were screened in the model group compared with the control group. Upregulated differential genes mainly include CCL3, CCL4, TNF, CXCL2, CCL5, CXCL10, TNFRsf1b, CD40, CCL2, IRGM1, BCL2, IFI204, IKB, IFNb1, GBP3, TNFaip3, GBP7, GBP5, GBP2, IL6, GBP2b, Fas, CD86, NOD2, IL12b, CD247, TRIM30d, CCR5, IL1bos, CXCL3, GNB4, CXCL9, PAK6, PYCARD, GNB3, PLCB1. As shown in figure 4B, 718 up-regulated genes and 1163 down-regulated genes were screened in the model group compared with the *kaji-ichigoside F1* intervention group. Upregulated differential genes mainly include CXCL2, CCL2, IL1b, TNFRsf1b, OAS1a, IFI204, IFNb1, BCL2, OAS1g, IRF7, IL6, STAT1, GBP5, GBP2, GBP3, GBP2b, IL18, GBP7, H2-T23, STAT2, H2-M3, CCL12, CD86, Fas, IL12b, TRIM30d, CD80, H2-Q2, CD247, RNAsel, CCR5, CXCL3, APLm2, CXCL9, GNG3, PLCB1.

**Figure 4.**
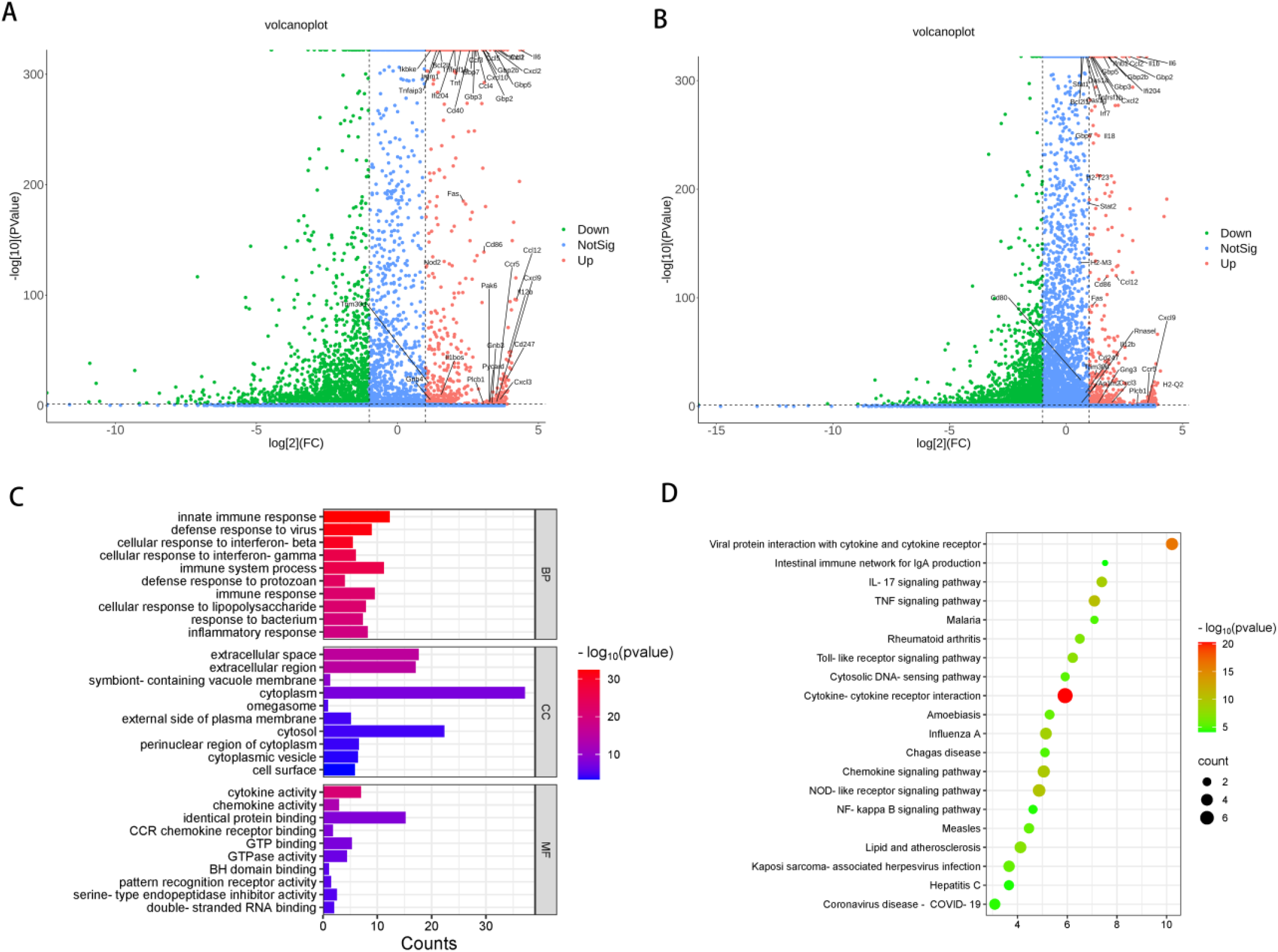
RNA sequencing results. A. up-regulated genes and down-regulated genes in the model group compared with the control group; B. up-regulated genes and down-regulated genes in the model group compared with the *kaji-ichigoside F1* intervention group; C. GO enrichment; D. KEGG enrichment.

#### 3.4.2. GO functional enrichment analysis

As shown in figure 4C, the up-regulated differences screened from the model group and the control group, as well as the up-regulated differential genes screened from the model group and the *kaji-ichigoside F1* intervention group were interleaved for further GO functional enrichment analysis: 1) In terms of BP, genes are significantly enriched in innate immune response, defense response to viruses, cellular response to interferon, immune response, cellular response to lipopolysaccharides, and inflammatory response to bacteria; 2) In terms of MF, differential genes were significantly enriched in cytokine activity, chemokine activity, CCR chemokine receptor binding, GTP binding, GTPase activity, BH domain, binding pattern recognition receptor, active serine endopeptide inhibitors, active double-stranded RNA binding and other functions; 3) In terms of CC, differential genes were significantly enriched in extracellular space, extracellular region, cytoplasm,extemal side of plasma membrane and cell surface.

#### 3.4.3. Enrichment analysis of KEGG signal pathway

As shown in figure 4D, the up-regulated differences screened from the model group and the control group, as well as the up-regulated differential genes screened from the model group and the *kaji-ichigoside F1* intervention group were interleaved for further KEGG pathway enrichment analysis, and the first 15 pathways were shown. It is mainly enriched in TNF signaling pathway, NOD-like receptor signaling pathway, IL-17 signaling pathway, TLR signaling pathway and NFKB signaling pathway, etc.

## Discussion

The present study integrates network pharmacology, transcriptomics, and molecular docking to elucidate the potential mechanism of *kaji-ichigoside F1*, a triterpenoid derived from *rosa roxburghii*, in ameliorating bacterial pneumonia. Our findings suggest that *kaji-ichigoside F1* modulates macrophage mediated inflammatory responses by targeting key signaling pathways, thereby reducing the pathological effects of *klebsiella pneumoniae* infection. Below, we discuss the implications of these results in the context of existing literature and highlight avenues for future research.

The network pharmacology analysis identified TLR4, NFKB1, STAT3, IL1B, and JUN as critical targets intersecting *kaji-ichigoside F1* and bacterial pneumonia. These genes are central to the regulation of inflammatory responses and macrophage polarization. TLR4, a pattern recognition receptor, is essential for detecting bacterial pathogens like *klebsiella pneumoniae* and initiating downstream pro-inflammatory signaling via NFKB and MAPK pathways^[12]^. The engagement of TLR triggers sequential molecular events culminating in the stimulation of nuclear regulatory proteins^[13]^. These activated transcription factors subsequently enhance the expression of multiple downstream genes responsible for producing immunomodulatory molecules, including intercellular signaling proteins, leukocyte attractants, mitogenic polypeptides, and biochemical modulators of inflammatory responses^[14]^. Our molecular docking results confirmed strong binding affinities between *kaji-ichigoside F1* and TLR4 (−7.6 kcal/mol), this aligns with studies showing that some natural compounds targeting TLR4 reduce excessive cytokine production in inflammatory models^[15]^. NFKB, a crucial regulatory protein in immune response mechanisms, exerts dual control over both innate and adaptive immunity while functioning as a central coordinator of inflammatory processes, was another key target. This transcriptional activator not only drives the synthesis of multiple inflammatory mediators through upregulating target genes but also directly modulates inflammasome activity at the molecular level^[16]^. The docking energy of *kaji-ichigoside F1* with NFKB1 (−7.4 kcal/mol) suggests that *kaji-ichigoside F1* may inhibit NFKB nuclear translocation, thereby attenuating pro-inflammatory gene expression. This mechanism is consistent with the observed downregulation of pro-inflammatory gene IL-1β, IL-6, and TNF-α mRNA in *kaji-ichigoside F1* intervention macrophages. Similar effects have been reported for ursolic acid, another triterpenoid, which suppresses NFKB, leads to anti-inflammatory effects^[17]^. STAT3, a mediator of M1/M2 macrophage polarization^[18]^, also emerged as a critical target, STAT3 activation promotes M1 polarization and sustains inflammation by amplifying cytokine signals^[19]^. The high binding affinity of *kaji-ichigoside F1* with STAT3 (−8.4 kcal/mol) implies its role in rebalancing macrophage phenotypes. This is supported by transcriptomic data showing *kaji-ichigoside F1* mediated downregulation of STAT1 and IL-12, which are markers of M1 polarization. Such modulation could shift macrophages toward an anti-inflammatory M2 phenotype, as evidenced by increased IL10 expression.

Macrophage polarization is pivotal in bacterial pneumonia progression^[20]^. Our result showed *klebsiella pneumoniae* infection skewed RAW264.7 cells toward an M1 phenotype, characterized by pseudopodia formation and pro-inflammatory cytokine secretion, such as IL-1β, IL-6, and TNF-α. *kaji-ichigoside F1* intervention restored cellular morphology, indicative of M2 polarization, it shows an increase in IL-10. This shift aligns with the observed increase in IL-10, a hallmark of M2 macrophages that resolves inflammation and promotes tissue repair ^[21]^. The modulation of STAT3 and JUN by *kaji-ichigoside F1* likely underpins this phenotypic switch, as both genes regulate macrophage plasticity^[22-23]^. RNA sequencing revealed that *klebsiella pneumoniae* infection upregulated genes associated with innate immunity (e.g., CCL3, CXCL2, TNF) and downregulated anti-inflammatory mediators. *kaji-ichigoside F1* intervention reversed these trends, suppressing pro-inflammatory chemokines (CCL2, CXCL3) while enhancing genes involved in immune resolution (IL-10).

KEGG enrichment highlighted the TLR, NFKB signaling pathways as central to *kaji-ichigoside F1* ‘s activity, corroborating network pharmacology predictions. TLR signaling pathways play critical regulatory roles in determining macrophage polarization states^[24]^. Experimental studies demonstrate that LPS, the canonical TLR4 ligand, serves as a potent driver of macrophage differentiation into the classically activated M1 subtype^[25]^. Molecular characterization reveals that LPS-mediated activation engages two distinct TLR signaling adapters-MyD88 and TRIF, which coordinate bifurcated downstream responses^[25]^. Substantial genetic evidence confirms the essential function of MyD88-dependent TLR cascades in establishing M1 polarization patterns and mediating inducible pro-inflammatory cytokine production^[26]^. Mechanistically, MyD88-driven signaling initiates IRAK kinase activation cascades, culminating in TRAF6 E3 ligase autoubiquitination and subsequent ubiquitin chain assembly on signaling intermediates required for TAK1 kinase activation^[25]^. Subsequently, TAK1 propagates signals through IKK complex activation, triggering IκBα phosphorylation and proteasomal degradation that ultimately liberates NFKB for nuclear translocation^[27]^. This transcriptionally active NFKB complex orchestrates the M1-specific inflammatory transcriptome by binding promoters of genes encoding TNF-α, IL-1β, IL-6, IL-12, and COX-2, thereby establishing its central regulatory position in macrophage polarization^[24]^. Notably, the IL-17 and NOD-like receptor pathways were also enriched, suggesting broader immunomodulatory effects^[28-29]^. IL-17 synergizes with TLR signaling to exacerbate neutrophil recruitment and tissue damage in pneumonia^[30]^, and *kaji-ichigoside F1*’s inhibition of this axis may further mitigate lung injury. The transcriptomic data also implicated interferon-related genes (IFNb1, IFN7) and GTPases (Gbp2, Gbp5) in *kaji-ichigoside F1*’s mechanism. These genes are critical for host defense against intracellular pathogens, but their overexpression during prolonged infection can drive cytokine storms^[31-32]^. *kaji-ichigoside F1*’s ability to temper their expression while maintaining antiviral responses reflects a balanced immunomodulatory profile, a desirable trait for treating bacterial pneumonia without compromising host defense.

## Limitations and Future Directions

While this study provides valuable insights, several limitations warrant attention. First, the analysis relied on *in vitro* macrophage models, which may not fully replicate the complexity of lung tissue microenvironment. Future studies should validate these findings in animal models of bacterial pneumonia. Second, the transcriptomic data focused on mRNA expression, proteomic analyses from animal lung tissue are needed to confirm translational effects. Third, the specificity of *kaji-ichigoside F1*’s interactions with TLR4, NFKB, STAT3 and JUN requires further validation using knockout models or siRNA silencing.

## Conclusion

In summary, this study demonstrates that *kaji-ichigoside F1* alleviates bacterial pneumonia by targeting TLR and NFKB signaling pathways, rebalancing macrophage polarization with suppressing pro-inflammatory cytokines and increasing anti-inflammatory cytokines. These findings position *kaji-ichigoside F1* as a promising candidate for developing adjunct therapies against bacterial infections. Future research should explore its pharmacokinetics, toxicity, and synergistic potential with existing antibiotics to translate these findings into clinical applications.

## Authors’contributions

Junlin Yang designed the research. Junlin Yang, Xi Yang, Min Cheng and Liping Wu performed the experiments and analysed the data. Junlin Yang wrote and revised the manuscript. All authors read and approved the final manuscript.

## Acknowledgements

This research was funded by the Guizhou Provincial Science and Technology Projects, China (grant number: [2024] No. Youth 282), the Guizhou Provincial Department of Education, China (grant number: [2024] No. 104) and the Science and Technology Fund project of Guizhou Provincial Health Commission, China (grant number: gzwkj2025-236).

## Ethics statement

The Ethics Committee of the Affiliated Hospital of Guizhou Medical University approved all procedures.

## Consent to participate

Not applicable

## Consent for publication

All authors gave their final approval and agreement to be accountable for all aspects of the work.

